# Disrupted Surfaces of Porous Membranes Reduce Nuclear YAP Localization and Enhance Adipogenesis through Morphological Changes

**DOI:** 10.1101/2021.01.31.429012

**Authors:** Zahra Allahyari, Stephanie M. Casillo, Spencer J. Perry, Ana P. Peredo, Shayan Gholizadeh, Thomas R. Gaborski

## Abstract

The disrupted surface of porous membranes, commonly used in tissue-chip and cellular co-culture systems, is known to weaken cell-substrate interactions. Here, we investigated whether disrupted surfaces of membranes with micron and sub-micron scale pores affect YAP localization and differentiation of adipose-derived stem cells (ADSCs). We found that these substrates reduce YAP nuclear localization through decreased cell spreading, consistent with reduced cell-substrate interactions, and in turn enhance adipogenesis, while decreasing osteogenesis.

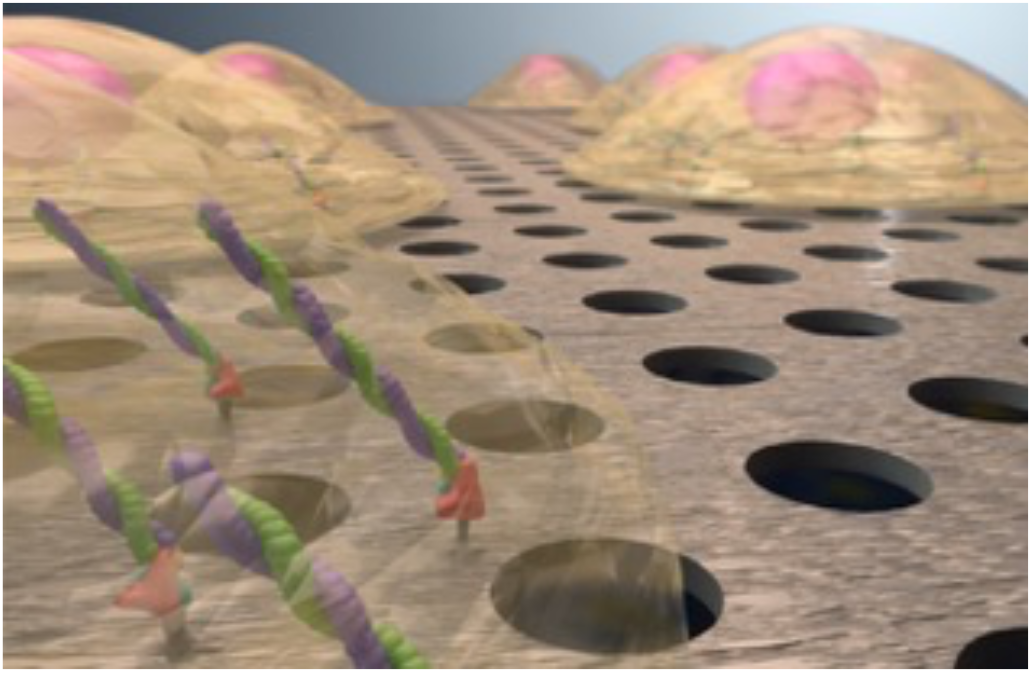

---

There is growing interest in disrupted substrates in the form of porous membranes, microposts, and micropillars in tissue engineering applications and stem cell studies due to their ability to tailor and recapitulate a physiologically relevant microenvironment.^1–10^ Porous membranes are possibly the most broadly applied disrupted surfaces in cellular studies and the rapidly growing field of tissue-on-a-chip.^2,11^ These membranes can enable physiologically relevant cell-cell communication between co-cultured cells and are widely used in developing organ-on-a-chip and barrier models.^1–6,12^ Although there have been several informative studies investigating the role of nano and micro topographies in cell-substrate interactions and mechanotransduction,^13–16^ the results from these studies are rarely applied or considered when using porous membranes in cell culture or tissue-on-a-chip devices. This issue is most apparent with microporous membrane applications where membranes are used as a semi-permeable transmigration barrier or to compartmentalize different cell types, but not to intentionally modify or perturb cell-substrate interactions. The role of surface discontinuity on cellular behavior of vascular endothelial cells was previously explored by our group, and it was demonstrated that membrane pores can alter fibronectin fibrillogenesis and cell migration, as well as F-actin polymerization, and focal adhesion formation.^11,17^ In another study, we showed that many of the observed changes in cellular behaviors over porous membranes could also be replicated by non-fouling micro-patterns which resemble membrane pores with regular disruption in cell-substrate interactions.^18^ These results suggest that surface disruption is responsible for the reduced or weakened cell-substrate interactions and downstream effects.

Mechanotransduction is the mechanism by which cells sense and respond to the mechanical forces and physical properties of their microenvironment. The effects of these mechanical cues can drive cell differentiation, determine cell fate, and define tissue architecture.^19^ Yes-associated protein (YAP) is a major transcriptional coactivator and a hippo pathway effector that mediates mechanotransduction while activated by localization to nuclei.^20,21^ YAP localization and activation are strongly correlated with substrate stiffness and associated cell-substrate interactions,^19,22–25^ and it has been shown that YAP can direct mesenchymal stem cell fate toward an osteogenic lineage on stiffer substrates.^22,26^ Although our previous efforts confirmed the formation of weaker vascular endothelial cell-substrate interactions over disrupted substrates similar to soft substrates, it has not been investigated whether these weaker cell-substrate interactions are correlated with altered mechanotransduction signaling pathways, and if they are associated with YAP nuclear localization and cell fate.

In this study, porous membranes with micron and submicron size pores and two different porosities were first used to explore how surface disruption might affect cell-substrate interactions and mechanotransduction in adipose-derived stem cells (ADSCs). The results indicate weakened cell-substrate interaction as well as reductions in YAP nuclear localization on all porous membranes, which is similar to the cellular behavior on soft substrates and tissues.^19,27,28^ We also confirmed that surface disruption on rigid non-fouling micropatterned substrate yielded similar and consistent results. Since there is an increasing interest in utilizing disrupted surfaces in mesenchymal stem cell differentiation, ^3,29–31^ we also evaluated whether the surface discontinuity of porous membranes could affect stem cell fate and the relationship to morphological changes in ADSCs.

Mesenchymal cells, including ADSCs, are established models in mechanoresponse and substrate stiffness-related studies.^32–34^ Here, we first sought to confirm whether ADSCs, like vascular endothelial cells, generate weaker cell-substrate interactions over discontinuous surfaces. The first step for understanding _surface_ disruption-related changes in cell adhesion and mechanotransduction required the fabrication of porous membranes with various pore sizes and porosities. Free-standing chip-supported silicon dioxide (SiO_2_) membranes with two different pore sizes (0.5 or 3 µm) and either low or high porosity (5 or 25 %) were fabricated using a combination of microfabrication techniques as reported in our previous work (Supporting Information).^35^ These two pore sizes were chosen since they are in the range widely used in various co-culture and cellular barrier studies.^2,18,36–38^ The pores were developed in the form of 0.5 µm-diameter circles with 1 µm or 2 µm center-to-center spacings and 3 µm-diameter circles with 6 µm or 12 µm center-to-center spacings in a hexagonal pattern (Figure 1), referred to as 0.5LP, 0.5HP, 3LP and 3HP, where LP refers to low porosity (5%) and HP is high porosity (25%).

**Figure 1.**
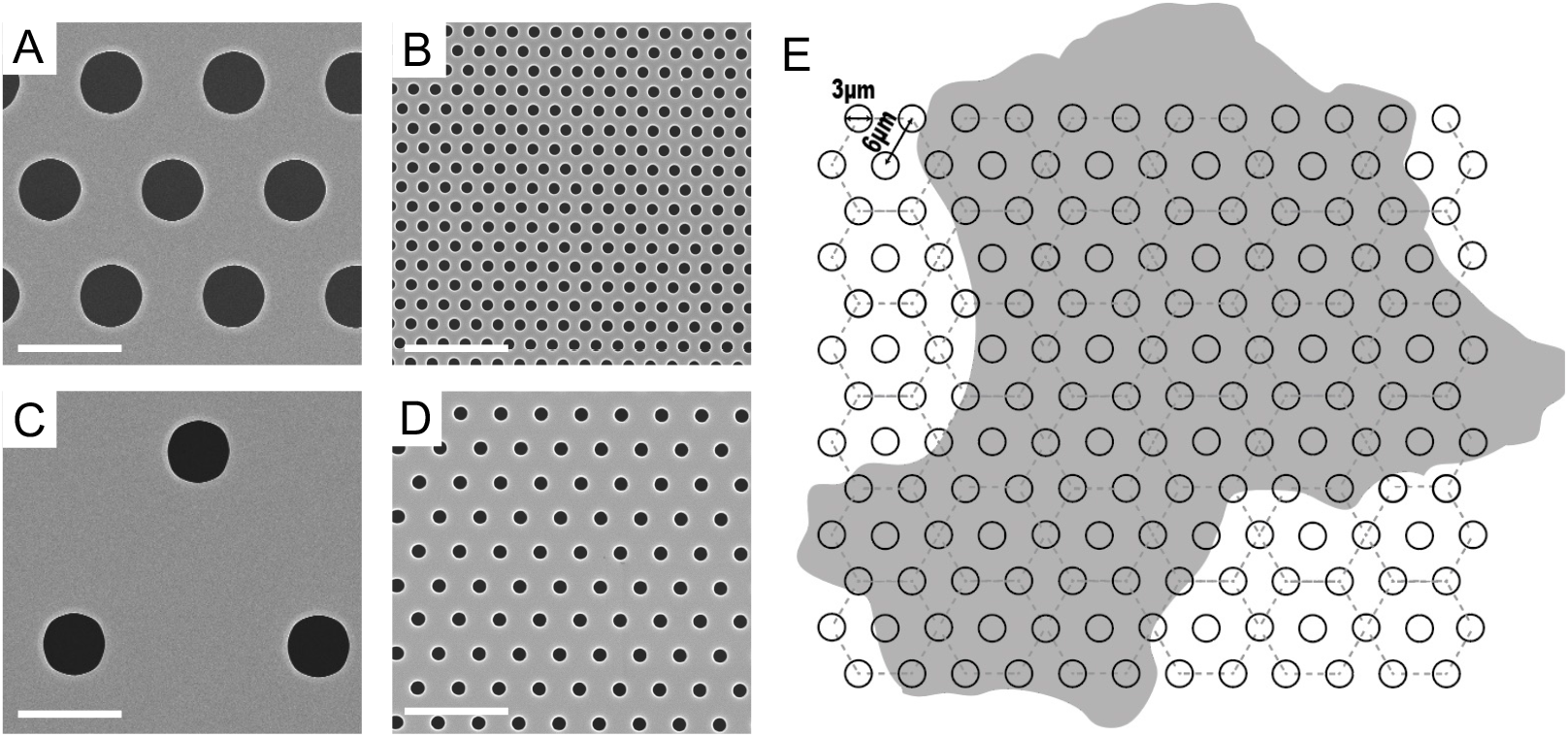
Identical magnification scanning electron micrographs of SiO_2_ membranes with (A) 3.0 µm diameter pores and high porosity, 3HP; (B) 0.5 µm diameter pores and high porosity, 0.5HP; (C) 3.0 µm diameter pores and low porosity, 3LP; and (D) 0.5 µm diameter pores and low porosity, 0.5LP. (E) Illustration of a characteristic cell outline on a 3.0 µm diameter pore and 25% porosity membrane, 3HP. White scale bars are 5 µm.

We first evaluated cell spreading, as an indicator of cell-substrate interaction strength^39^ over porous membranes and non-porous controls after 24 h of cell culture. Up to 30 percent reduction was observed in cell spreading over the porous membranes compared to the non-porous SiO_2_ substrate (Figure 2). Not surprisingly, we also found significantly reduced F-actin in cells cultured over the porous membranes, even when normalized for cell area (Table S1). The number of focal adhesions was not significantly different between substrates (Table S1). Reduced cell spreading over porous membranes may be due to the diminished cytoskeleton formation and contractility as a result of weaker cell-substrate interaction. To test this hypothesis, ADSCs were treated with 1µM Cytochalasin D (CytoD) for 24 h. CytoD is an inhibitor of actin dynamics, disrupting actin polymerization and network organization. CytoD-treated ADSCs demonstrated significantly smaller spreading areas over non-porous membranes compared to untreated cells. However, cell spreading remained almost the same after CytoD treatment on the porous membranes. Interestingly, treated cells on non-porous membranes had similar spreading to untreated cells on porous membranes.

**Figure 2.**
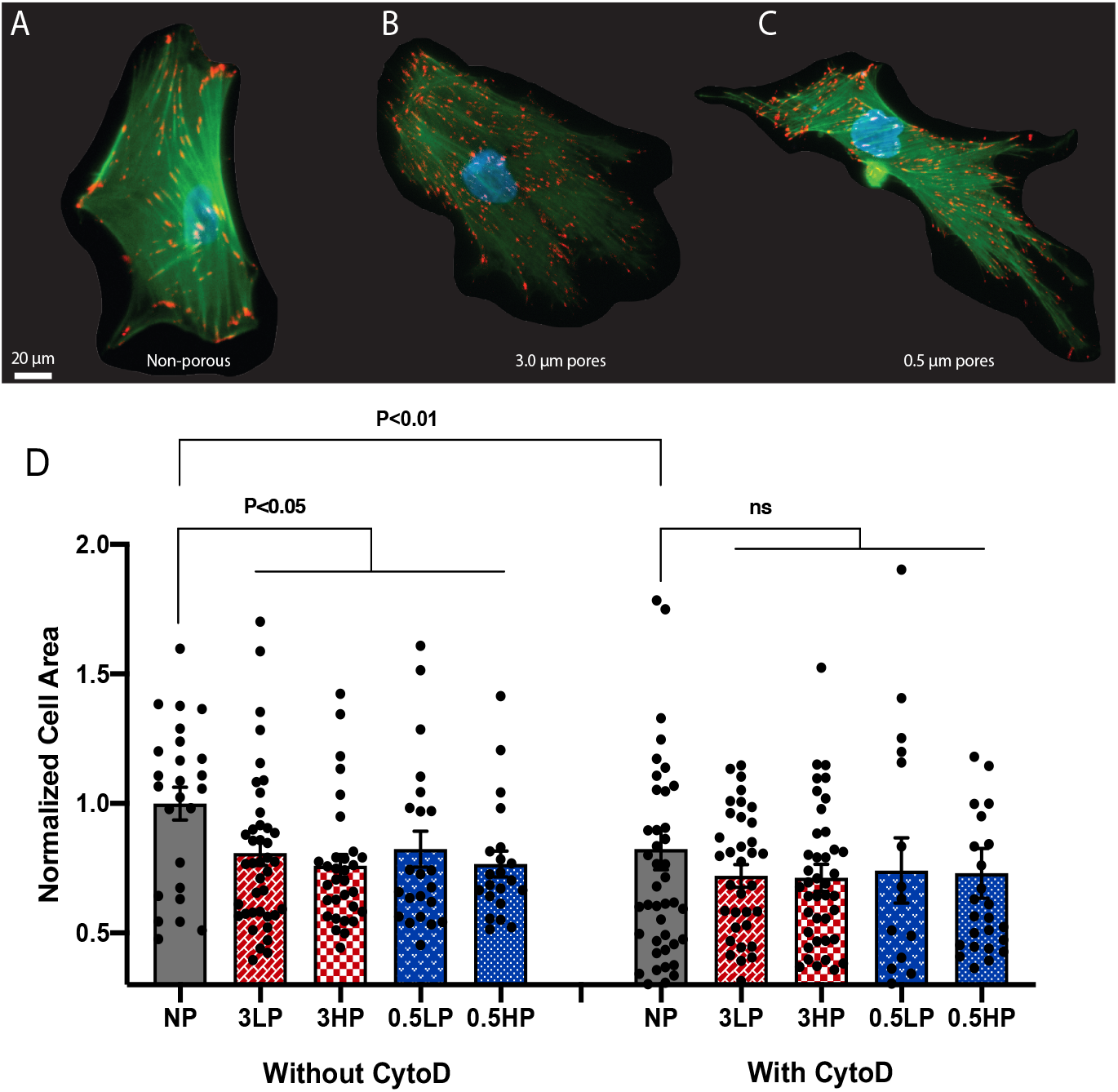
Fluorescence images of ADSCs stained for F-actin (phalloidin, green), focal adhesions (anti-vinculin, red) and nuclei (DAPI, blue) on (A) Control non-porous SiO_2_ membrane (NP), (B) 3.0 µm pores SiO_2_ membrane, and (C) 0.5 µm pores SiO_2_ membrane. (D) Cell spread area over various membranes after 24h of cell seeding with and without CytoD. (Cell areas are normalized to the mean cell area on nonporous controls.) (n = 4 substrates).

We then aimed to investigate mechanotransduction of ADSCs cultured over porous membranes using YAP localization as an indicator. The YAP expression was assessed by immunofluorescence staining 6 h and 24 h after cell seeding (Figure 3). The nuclear and perinuclear YAP content was measured in the captured images to calculate the ratio of nuclear to cytoplasmic YAP localization. We looked at YAP localization 6 h after cell seeding and found no significant difference between the control and porous membranes, which was not surprising as the cells were still adhering to and spreading on the substrate. After 24 h, however, we observed a statistically significant difference in nuclear YAP localization over the discontinuous porous membranes versus the continuous non-porous control (Figure 3F). The results indicated that YAP entered the nuclei after ADSCs sensed the stiffness of the rigid continuous SiO_2_ substrate. However, the cells did not behave similarly on the porous membranes, and despite the fact that ADSCs were placed on a rigid substrate, the nuclear to cytoplasmic YAP ratio was significantly less over all the porous membranes. To determine the relationship between diminished cell spreading resulting from CytoD treatment and nuclear YAP reduction, the YAP localization of CytoD-treated ADSCs were evaluated on porous membranes and non-porous substrates. The YAP nuclear localization of CytoD-treated ADSCs on the non-porous substrate decreased to the same level of the YAP localization on the porous membranes. These observations demonstrate that the surface disruption and weakened cell-substrate attachment can affect mechanotransduction despite the substrate being extremely stiff (Young’s modulus >100 GPa). Additionally, these results are consistent with Nardone et al.’s finding where they showed that YAP nuclear localization can be induced independently of FA formation.^21^ They suggested that although YAP can direct FA-related gene expression, FA assembly does not necessarily lead to YAP nuclear localization. It was shown that cell spreading rather than FA assembly regulated YAP localization.^21^ Here, we also concluded that cell spreading rather than FA assembly plays the main role in mechanosensing and YAP localization, since we observed a significant decrease in nuclear YAP localization without a significant reduction in FA formation (Table S1).

**Figure 3.**
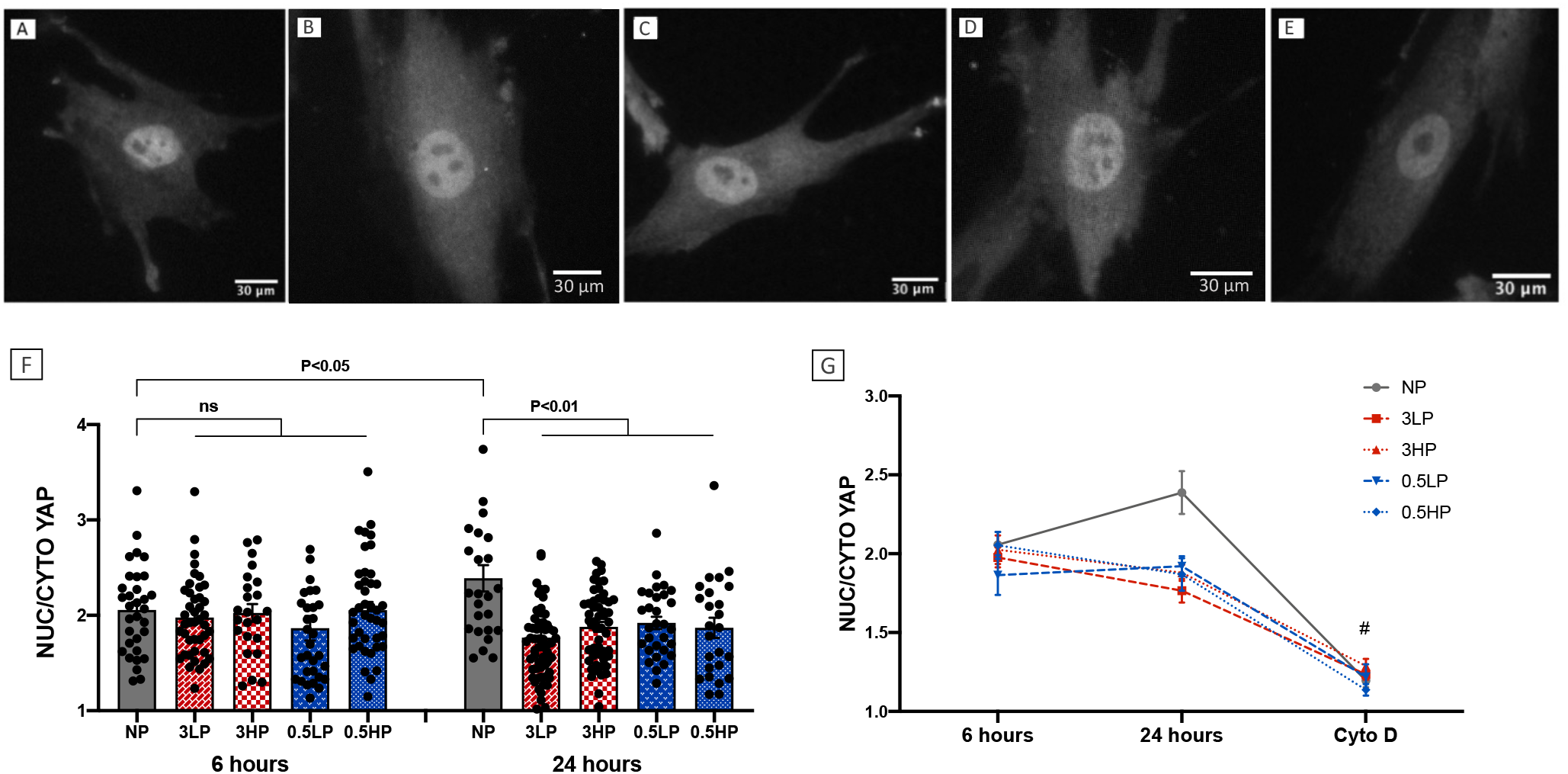
Fluorescence images ADSCs stained with anti-YAP on (A) Control non-porous SiO_2_ membrane, (B) 3.0 µm diameter pores and ∼5% porosity, (C) 3.0 µm diameter pores and ∼25% porosity, (D) 0.5 µm diameter pores and ∼5% porosity, and (D) 0.5 µm diameter pores and ∼25% porosity. (F) YAP nuclear-cytoplasmic ratio is quantified at 6 h and 24 h. (G) Changes in YAP ratios from 6h to 24h and after exposure to 1 µM CytoD at 24 h. # Cells on non-porous substrate saw a significant decrease in YAP after exposure to CytoD and reached to the same level of the YAP localization on porous membranes.

We further aimed to confirm the role of surface disruption on mechanotransduction by repeating the experiment on a non-fouling micropatterned substrate. We developed a non-fouling micropatterned surface with an identical hexagonal pattern to membranes’ pores and the same SiO_2_ surface chemistry (Supporting Information). Briefly, the non-fouling regions were generated in the form of 3 µm-diameter circles using poly(l-lysine)-grafted-poly(ethylene glycol) (PLL-g-PEG) on a silicon dioxide (SiO_2_) layer by combining photolithography and simple polymer adsorption on the surface, described in detail in our previous work. We seeded ADSCs on the developed PEG micropatterned and unpatterned substrates 6 and 24 hours before immunofluorescence staining and quantified cell spreading and YAP localization from the captured images. We found the same trend of reduction in cell spreading and nuclear YAP localization on the PEG micropatterned substrates versus unpatterned substrates as compared to the porous membranes versus non-porous substrates (Figure S2). Together, these results show that discontinuous planar surfaces affect mechanotransduction similarly to what has been previously shown on soft substrates.

Since there is a growing interest in the application of disrupted surfaces such as porous membranes in co-culture systems for differentiation studies due to their ability to compartmentalize cell culture, ^3,29,40–42^ we sought to understand whether the altered mechanotransduction over porous membranes could affect cell fate. We aimed to investigate ADSC differentiation into adipocyte and osteocyte lineages on porous and non-porous SiO_2_ membranes. ADSCs were seeded on the membranes and fed a 1:1 mixture of adipogenic and osteogenic media for 14 days similar to Guvendiren et al.^43^ Since there was no significant difference between spreading and mean YAP ratio on all four porous membranes, we chose high porosity 0.5 µm pore membranes (0.5HP) for the differentiation studies and histological staining and imaging, due to their better imaging properties in brightfield.

Cell morphology has long been studied and is known to be a significant contributing factor that can guide cell differentiation.^44–47^ Differences in contractility of cell cytoskeleton resulting from cell morphology can regulate mechanotransduction, which subsequently can direct cell fate.^44,48^ Cell elongation is one of the major studied morphological aspects of cells and can direct the cells towards osteogenesis whereas round morphology improves adipogenesis.^47,49^ For instance, Kilian et al. showed that shape cues with higher aspect ratios led to increased osteogenesis rather than adipogenesis in mesenchymal cells with the same adhesion area.^48^ Therefore, we decided to evaluate cell elongation following cell seeding on the membranes to determine the potential relationship between morphology and differentiation. We measured cell elongation (aspect ratio) of ADSCs 1, 7, and 14 days after seeding during the two-week bi-differentiation protocol (Figure 4). We hypothesized that the relatively stronger cell-substrate interactions over non-porous membranes would lead to more elongation, whereas cells over porous membranes might remain more rounded and less spread, consistent with the results in Figure 2D. We found that ADCSs cultured on porous versus non-porous membranes showed a slight but insignificant difference in elongation on day 1. By day 7 and through day 14 of the bi-differentiation protocol, we found cells on non-porous substrates were significantly more elongated. Interestingly, ADSCs with CytoD for all 14 days (++) showed similar elongation on both non-porous and porous membranes. However, removing CytoD from cell culture media after day 7 (+-) led to a rebound in elongation over non-porous membranes, while cells remained rounded on the porous membrane even after removing CytoD. This partially reversed morphology change on the non-porous substrate but not the porous membrane shows that CytoD effect is reversable and the cells are still responsive to the microenvironmental substrate cues.

**Figure 4.**
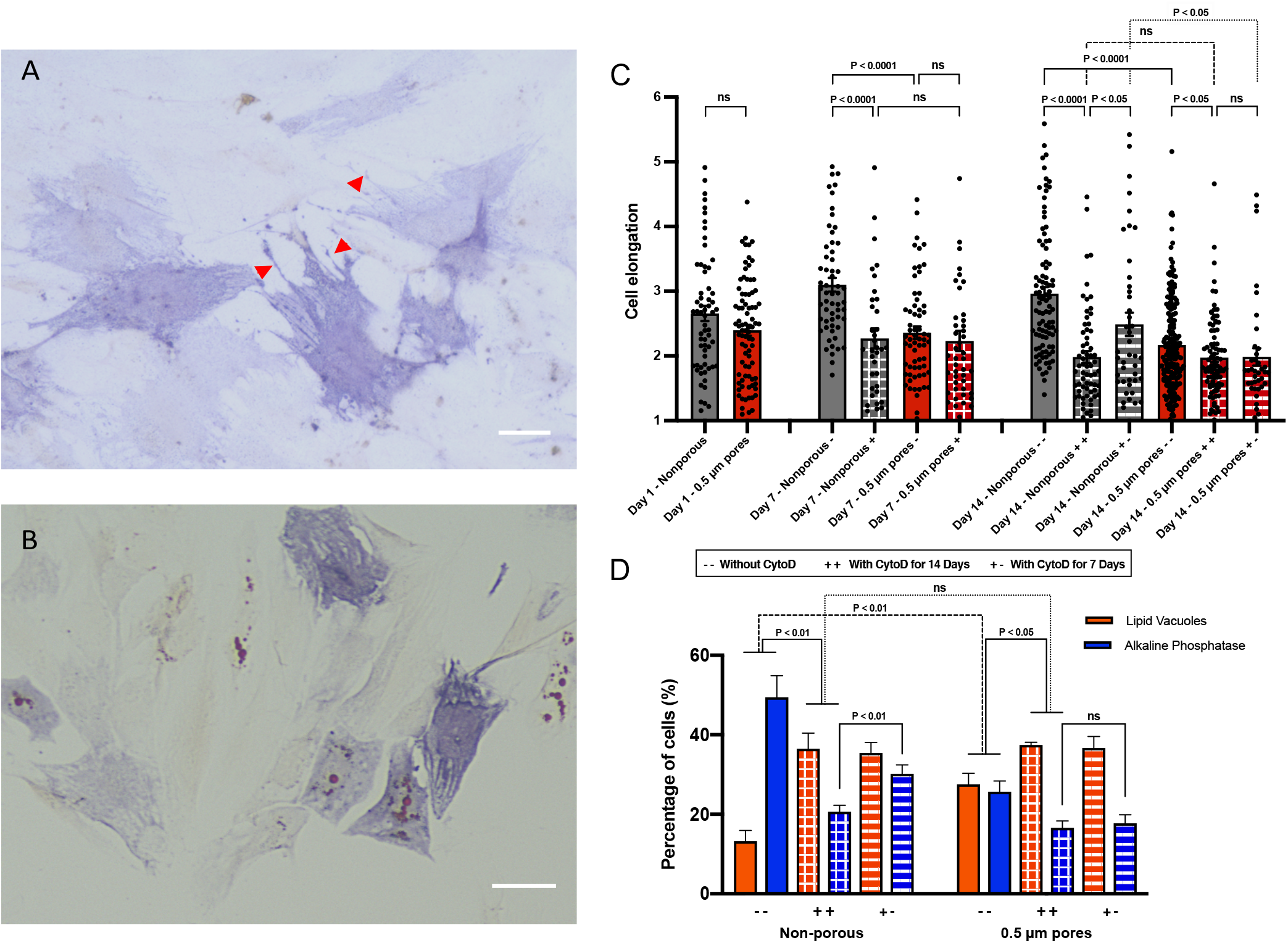
ADSCs were cultured in mixed adipogenic and osteogenic differentiation media on each substrate for 14 days and then prepared for histological staining. ALP and lipid vacuoles were respectively stained by Fast blue RR and Oil red O on (A) non-porous and (B) 0.5 µm pore diameter SiO_2_ membranes (Scale bars = 30 µm). Red Arrows on (A) point to some of the long, narrow filopodia on non-porous SiO_2_ which are mostly missing on (B) porous membranes. (C) Cell elongation of ADSCs 1, 7, and 14 days after cell seeding on non-porous and 0.5 µm pore diameter SiO_2_ membranes without CytoD (--), with CytoD for 14 days (++), and with CytoD for the first 7 days (+-) (n > 3 substrates for each condition; > 40 cells). D) percentage of the cells expressing lipid vacuoles and alkaline phosphatase which are stained by Oil red O and fast blue RR respectively on day 14 (n = 4 substrates for each condition; > 400 cells).

Another major morphological difference between ADSCs over these two substrates was the nature of cytoplasmic protrusions formed by the cells. While ADSCs formed several long, narrow filopodia on non-porous substrates, the dominant cytoplasmic protrusions on porous membranes were short, wide lamellipodia-like projections (Figure 4A, B). Filopodia were rarely found in ADSCs on porous membranes, even in cells that showed alkaline phosphatase (ALP) staining. This finding is consistent with previous reports where the formation of short, cytoplasmic protrusions was attributed to weak cell-substrate adhesion.^50^ Considering the known role of cytoplasmic protrusions in regulating mechanical signal transductions,^46^ the absence of these structures may be an indicator or contributing factor to cell fate over these porous membranes.

Analysis of the images stained for lipid vacuoles and ALP on day 14 confirmed our expectations that the differences in the earlier cell-substrate interaction experiments over non-porous and porous membranes would carry over to differentiation. We found that cells were more prone to generate lipid vacuoles, indicating adipogenic differentiation,^43,51^ over the disrupted substrates of porous membranes (Figure 4B). On the other hand, far fewer ADSCs expressed lipid vacuoles on the non-porous membranes at the same time point (Figure 4A). Instead, the cells over the non-porous samples showed more ALP staining than the cells cultured on porous membranes. Since ALP is an enzyme that is highly expressed in the early stage of osteogenesis,^52,53^ the enhanced ALP expression in the ADSCs on non-porous membranes shows a preference towards osteogenic differentiation (additional images are available in the Supporting Information file, Figure S3). This might be expected based on the morphology data as osteoblasts normally present more elongated forms with higher contractile cytoskeleton as compared to adipocytes. Interestingly, these distinct morphologies were readily observed by day 7 of the two-week differentiation protocol when differentiation markers were not yet substantially expressed (Figure 4C). This finding suggests that the lack of strong cell adhesion required for the formation of contractile cytoskeleton might hinder the formation of more elongated morphology in the cells over porous membranes and subsequently contribute to regulating mechanotransduction and directing cell fate in addition to the role of cell spreading. To confirm this idea, we also evaluated cell differentiation of ADSCs treated for all 14 or just the first 7 days with CytoD. CytoD treatment for the differentiation period led to the increased adipogenic differentiation and decreased osteogenic differentiation on both porous and non-porous substrates, consistent with the earlier studies which demonstrated CytoD contribution in adipogenic differentiation^54,55^ However, differentiation changes were sharper on non-porous substrates, which removed the difference between differentiation over the two substrates. Interestingly, removing CytoD from differentiation media after day 7 improved osteogenic differentiate on non-porous substrates but not on the porous membranes. Adipogenic differentiation remained the same on the both substrates. It seems that some of the undifferentiated cells could repair their actin network and regain elongated morphology required for osteogenesis on non-porous membrane via generating effective cell-substrate interactions after removing CytoD. However, ADSCs were unable to improve their cytoskeleton assembly on porous membranes after stopping CytoD treatment, potentially due to the lack of strong cell-substrate interactions. Collectively, these results confirm that ADSC fate is altered on the discontinuous substrates, with a promotion of adipogenesis, similarly to the enhancement on soft substrates.^56^

Throughout our YAP and morphology analyses, we found statistically significant differences in mean values between cells on continuous versus disrupted surfaces. However, the plots showed a substantial spread and overlap in data points of individual cells between the two conditions. While this spread in data is not surprising for cellular studies, we were curious if the amount of overlap or non-overlap might be meaningful with regards to the percentage of cells that ultimately showed lipid vacuoles or were ALP positive after 14 days of the bi-differentiation protocol. We found that more than three times as many cells cultured on non-porous membranes were ALP positive versus presenting lipid vacuoles, whereas cells cultured over porous membranes showed a nearly equal distribution (Figure 4D). Interestingly, we noticed a similar proportion of cells above or below an arbitrary YAP ratio of 2. More than three times as many cells had a YAP ratio above 2 on non-porous membranes. On the other hand, a YAP ratio of 2 nearly equally split the data points on porous membranes (Figure 3E). The same proportions and YAP ratio is also true for the analyses on continuous and micropatterned PEG substrates in Figure 2. Analyzing the cell elongation data on day 14 in Figure 4C yet again shows the same proportions if a cell aspect ratio of just over two is selected – more than three quarters of the cells on non-porous substrates have a cell aspect ratio above two, while an aspect ratio of two nearly equally splits the data points on porous membranes. These correlating distributions warrant future studies to investigate if morphological and YAP measurements can predict an individual cell’s differentiation outcome, and whether some cells within a population are more influenced by disrupted and discontinuous surfaces.

In conclusion, these experiments demonstrate that substrate discontinuity, including that arising from porous membranes, affects mechanotransduction and cell differentiation similarly to soft substrates. ADSCs on discontinuous substrates produce different responses compared to continuous substrates, including weakened cell-substrate interaction, as evidenced by decreased cell spreading as well as reduced nuclear YAP localization. We later confirmed that altered mechanotransduction could lead to different cell fates as a result of the disrupted surfaces and morphological changes. Our findings demonstrate that membranes, commonly used in co-culture and tissue-chip applications, have the potential to significantly affect cell-substrate interactions including substrate-directed differentiation. The outcome of this study can be employed to better understand mechanotransduction and cell differentiation in barrier models and tissue-chips, and to design membranes that can encourage stem cell differentiation toward desired phenotypes.

## ACKNOWLEDGEMENTS

We thank Bradley Kwarta for producing the cell-substrate interaction illustrations. The authors declare the following competing financial interest: TRG is a co-founder of SiMPore and has an equity interest in this early-stage company commercializing ultrathin silicon-based technologies.

## AUTHOR CONTACT INFORMATION

Thomas R. Gaborski; twitter: @tomgaborski; ORCID: 0000-0002-3676-3208

## AUTHOR CONTRIBUTIONS

The manuscript was written through contributions of all authors. All authors have given approval to the final version of the manuscript.

## SUPPLEMENTARY INFORMATION

### METHODS AND MATERIALS

#### Fabrication of Ultrathin SiO_2_ Membranes

Free-standing chip-supported SiO_2_ membranes were fabricated using a combination of different microfabrication techniques as reported in our previous work.^1^ Tetra-ethoxysilane (TEOS)-based PE-CVD was used to deposit a SiO_2_ layer (300 nm) on both sides of a double-side-polished silicon wafer. The front-side was patterned with ASML PAS 5500/200 iline stepper to obtain hexagonally packed 0.5 μm and 3 μm openings (1 μm and 6 μm center-center spacing, respectively), and it was thoroughly etched with reactive ion etching (RIE) using Drytek 482 Quad Etching tool. This was followed by annealing in nitrogen at 600 °C for film stress stabilization and increasing film robustness. The back-side SiO_2_ layer was patterned and etched and the silicon wafer was etched with ethylenediamine pyrocatechol (EDP) to obtain final 5.4 mm × 5.4 mm square chips with 2 mm × 2 mm open windows, and the chips were cleaved for making individual chips. Similar to the non-fouling patterned substrates, silicone gaskets were fabricated and attached to the membranes to retain the cells on the membranes.

#### Generation of Non-fouling PEG Micropatterns on Silicon Dioxide (SiO_2_)

SiO_2_ substrates were fabricated and patterned using microfabrication techniques as previously reported.^2^ Briefly, SiO_2_ layer (300 nm) was deposited using plasma-enhanced chemical vapor deposition (PE-CVD) on a 150 mm single-side-polished silicon wafer. Upon applying MicroPrime MP-P20 as an adhesion promoter agent, Microposit S1813 photoresist was spin-coated, and soft-baked at 115 °C. This was followed by G-line exposure using GCA 6000-Series DSW 5X stepper to create 3 µm openings in the photoresist layer with hexagonally packed layout with 6 µm center-to-center spacing. The resist was developed with MF CD-26 developer and washed with DI water. The wafer was hard baked for 1 min at 150 °C. For development of non-fouling regions, 0.5 mg/ml of PLL(20)-g(3.5)-PEG(2) (Nanosoft Biotechnology LLC) in 10mM HEPES was placed on the substrates for 75 min. The optimum concentration and coating time were determined in an earlier work.^2^ The substrates were washed twice immediately after removing PLL-g-PEG solution, followed by acetone wash to remove the photoresist and rinsed again with DI water. The non-fouling characteristic of the coated regions was confirmed by preventing protein adsorption and cell adhesion in an earlier study.^2^ Silicone sheets were cut with pre-designed shapes with a digital craft cutter to obtain silicone gaskets for fabrication of cell culture devices. The chips were bonded to silicone gaskets facilitated by surface treatment with corona discharge using corona treater (Nbond, Littleton, CO) to restrain cell culture area.

#### Cell culture

Adipose-derived stem cells (ADSCs) were purchased from ThermoFisher Scientific (Waltham, MA), and cultured in MesenPRO RS Medium with 2% Mesen-Pro Growth Supplement, 1% L-glutamine, and 1% Penicillin-Streptomycin. All the cell culture reagents were purchased from ThermoFisher Scientific unless stated otherwise. ADSCs were detached using TrypLE and seeded in the density of 2300 cells/cm for differentiation tests and 6000 cells/cm for the rest of the tests on all samples including non-fouling patterned SiO_2_, porous membranes and control samples. ADSCs were used between passages 3-6.

#### Immunofluorescence Staining

After 24 h of cell culture, the cells permeabilized using 0.1% Triton X-100 for 3 seconds, and fixed with 3.7% formaldehyde for 15 min. ADSCs were blocked with 2% (20 mg/ml) BSA for 15 min, and stained with DAPI (300 nM) for 3 min, 1:400 AlexaFluor 488 conjugated phalloidin for 15 min, and 1:100 eFluor570 conjugated anti-vinculin, Clone 759 for 2 h in room temperature to visualize nuclei, F-actins and focal adhesions, respectively. All the aforementioned steps followed by triple washing with PBS or DI water. For imaging the cells on non-fouling patterned substrates, the cells were washed and 10 µL PBS was placed on each sample. The samples were flipped on coverslips to enable imaging on them. However, for imaging on the porous membranes, the cells washed and the samples were only flooded with PBS without flipping since it was only necessary for the non-transparent substrates. To visualize nuclei, F-actin and focal adhesions, the fluorescent images were acquired through DAPI, GFP and Texas red filters, respectively. For YAP localization, ADSCs were fixed with 3.7% formaldehyde for 15 min, followed by permeabilizing with 0.1% Triton X-100 for 3 min. The cells were blocked with 4% (40 mg/ml) BSA for 15 min and stained with 1:100 AlexaFluor 488 conjugated YAP for 2 h to visualize YAP content. All the mentioned steps followed by triple PBS washing. The stained samples with AlexaFluor 488 conjugated YAP were imaged using GFP filter. All images were captured using Keyence BZ-X700 microscope (Keyence Corp. of America, MA) or Leica DMI6000 microscope (Leica Microsystems, Buffalo Grove, IL).

#### Actin Stress Fiber and focal adhesion analysis

The mean intensity of the actin fibers and the background intensity were determined from the actin cytoskeleton-stained images in ImageJ software. This value multiplied by the spreading area of the cells to calculate the total F-actin content of each cell. Acquired fluorescent images of the stained cells with eFluor570 conjugated anti-vinculin were used to evaluate focal adhesions. Only cells with less than 10 percent cell-cell contact were analyzed. The number of distinct focal adhesions were counted for each cell.

#### Spreading Area and Cell Morphology

The spreading area of the cells was obtained from the actin cytoskeleton-stained images. The cell areas were calculated by identifying the perimeter of the cells manually using ImageJ software. DAPI images from nuclei confirmed each measured area belongs to a single cell, and the cell did not undergo mitosis. Cell elongation was measured as the ratio of the cell length to the cell width in the fluorescent stained cells. The cell’s longest axis was defined as its length, and the longest axis perpendicular to the length was determined as its width. To inhibit actin polymerization and cytoskeletal contractility, cells were treated with 1 μM cytochalasin D in dimethyl sulfoxide (DMSO) at each medium change. Control cells were treated with DMSO as the vehicle control.^3^

#### YAP nuclear localization

The fluorescent images of the stained cells using AlexaFluor 488 conjugated YAP were utilized to analyze YAP nuclear localization. The freehand selection tool of ImageJ software was used to measure the mean intensity value in the nucleus, in the cytoplasmic perinuclear region less than 10 um away from the nucleus, and in the background immediately adjacent to the cell. The background intensity was subtracted from the nuclear and cytoplasmic intensity values for each cell. In order to obtain the YAP nuclear localization ratio, the background-subtracted nuclear intensity was divided by the background-subtracted cytoplasmic intensity.

#### Adipogenic and Osteogenic Differentiation

ADSCs were seeded in the density of 2300 cell/cm2 on the 0.5 µm and nan-porous control SiO_2_ in MesenPRO RS Medium. The MesenPRO RS Medium was removed, and a half and half mixture of adipogenic and osteogenic media with 1 % pen/strep were added to the samples after 24 h. ADSCs were cultured in this mixture for 14 days, and the media were changed every 3 days. ADSCs were fixed in 3.7% formaldehyde for 1 min and washed with DI water. The samples were stained for alkaline phosphatase and lipid vacuoles using Fast Blue RR salt (Sigma-Aldrich) and Oil Red O (Sigma-Aldrich) to evaluate their differentiation over each sample. Briefly, a pre-weighted capsule of fast blue RR salt was dissolved in 48 ml DI water, and 2 ml Naphthol (Sigma-Aldrich) was added to the solution. The solution was added to the fixed samples and washed twice with DI water after 30 min. Cells were incubated in 60% isopropanol, and the solution was removed after 5 min. Oil Red O working solution was prepared by adding 2 parts water to 3 parts 0.5 % Oil Red O in isopropanol. ADCSs were incubated in the Oil Red O working solution for 30 min and washed with DI water 4 times. The nuclei of ADSCs were stained using DAPI to assist with cell counting and calculating the percentage of the cells which went through osteogenic or adipogenic differentiation. Brightfield images were captured from each sample to visualize their lipid vacuoles and alkaline phosphatase. The differentiation on each sample was calculated using the overlap of the brightfield and DAPI fluorescent images.

#### Statistical Analysis

For statistical analysis, data with Gaussian distributions were evaluated with two-tailed student’s t-test, and Mann−Whitney test was used for the rest of the data. Sample sizes are noted in each Figure legend.

### Cellular F-actin and Focal Adhesions over Porous Membranes and Micropatterned PEG Substrates

Since actin stress fibers are ubiquitous and essential for cell adhesion and mechanosensing,^2,4–6^ F-actin content of the ADSCs was evaluated after 24 h of cell seeding (Figure S1A, B). The cells showed significantly lower F-actin content on the discontinuous surface of the Porous Membrane and non-fouling patterned SiO_2_ substrate compared to the continuous SiO_2_ (Figure S1C). The diminished F-actin formation on the disrupted substrate suggests weakened interactions of ADSCs with the substrates similarly to endothelial cells on porous membranes and PEG micropatterned substrates in our previous studies.^2,7,8^ In the next step, we evaluated FA formation as the most studied and largest adhesion structures in the cells.^9,10^ FAs are known to be key players in mechanosensing, and they are considered a bridge between substrate and cytoskeleton.^11–13^ FA formation was quantified by measuring the number of distinct focal adhesions. There was a slight but insignificant decrease in the number of distinct FAs and they were smaller and less elongated over the micropatterned substrates (Figure S1D).

**Table S1.** F-actin intensity per cell and distinct focal adhesions per cell on the non-porous substrate, porous membranes with 3 µm or 0.5 µm diameter pores and 3 µm PEG islands. (* F-actin formation was significantly different on discontinues substrates compared to continuous SiO_2_, but the number of focal adhesions was not significantly different on all samples)

| Substrate | F-actin intensity per cell ($\times 10^8$) | Number of focal adhesions per cell |
| --- | --- | --- |
| Continuous SiO<sub>2</sub> | $3.656 \pm 0.551$ | $309.5 \pm 72$ |
| 3 $\mu\text{m}$ diameter pores | $1.315 \pm 0.246$ | $236.3 \pm 57.6$ |
| 0.5 $\mu\text{m}$ diameter pores | $0.836 \pm 0.115$ | $251.5 \pm 53.5$ |
| 3 $\mu\text{m}$ diameter PEG islands | $1.871 \pm 0.241$ | $254 \pm 76$ |

**Figure S1.**
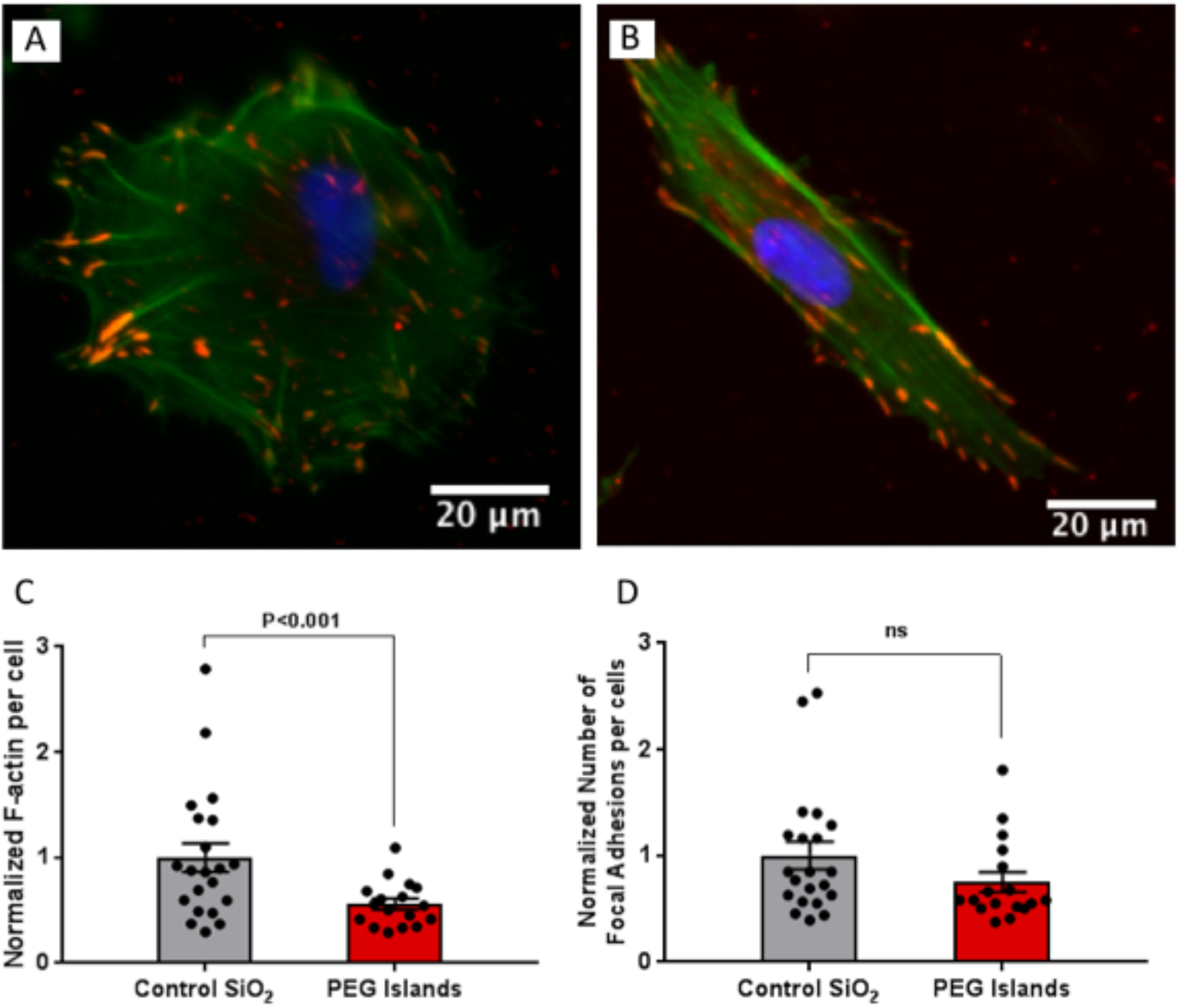
Fluorescence images of ADSCs stained for F-actin (phalloidin, green), focal adhesions (anti-vinculin, red) and nuclei (DAPI, blue) on (A) Control unpatterned SiO_2_ substrates and (B) SiO_2_ substrates with PEG islands after 24 h. (C) F-actin intensity per cell normalized to control mean value. (D) Distinct focal adhesions per cell normalized to control mean value.

### Cell Spreading and YAP Nuclear Localization on Non-fouling Micropatterned Substrate

To confirm the role of surface disruption on mechanotransduction, cell spreading and nuclear YAP localization were evaluated non-fouling micropatterned substrate with the same pattern of surface disruption as 3.0 µm diameter pores and ∼25% porosity. The same trend of reduction in cell spreading and nuclear YAP localization were observed on the PEG micropatterned substrates compared to unpatterned substrates which showed that disrupted surfaces of rigid substrates could affect mechanotransduction similarly to soft substrates.

**Figure S2.**
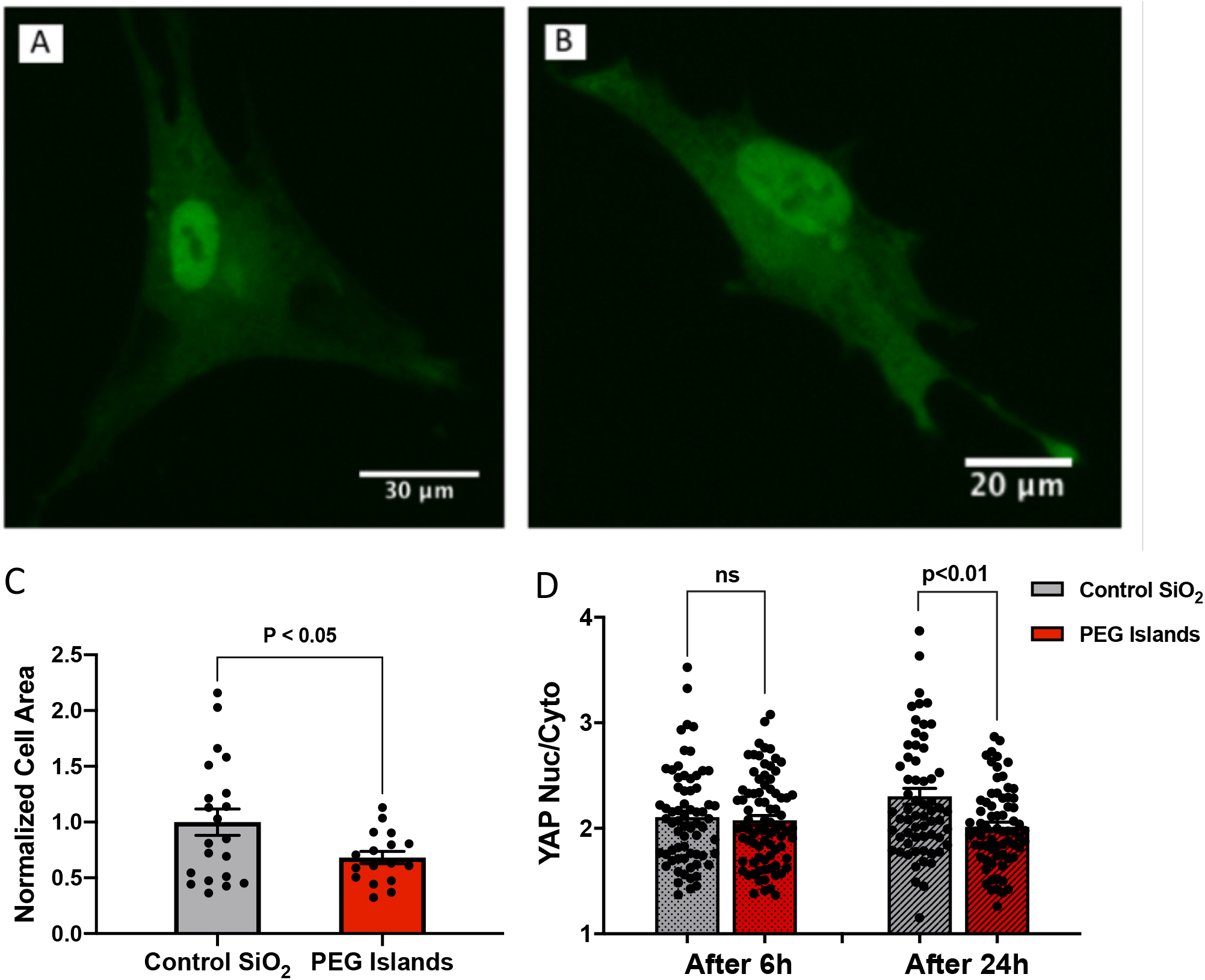
Fluorescence images of ADSCs stained with anti-YAP on (A) Control unpatterned SiO_2_ substrates and (B) SiO_2_ substrates with PEG islands. (C) Cell spread area was quantified from F-actin images. (D) YAP nuclear-cytoplasmic ratio is quantified at 6 h and 24 h after cell seeding (n = 4 substrates for each condition; > 50 cells).

**Figure S3.**
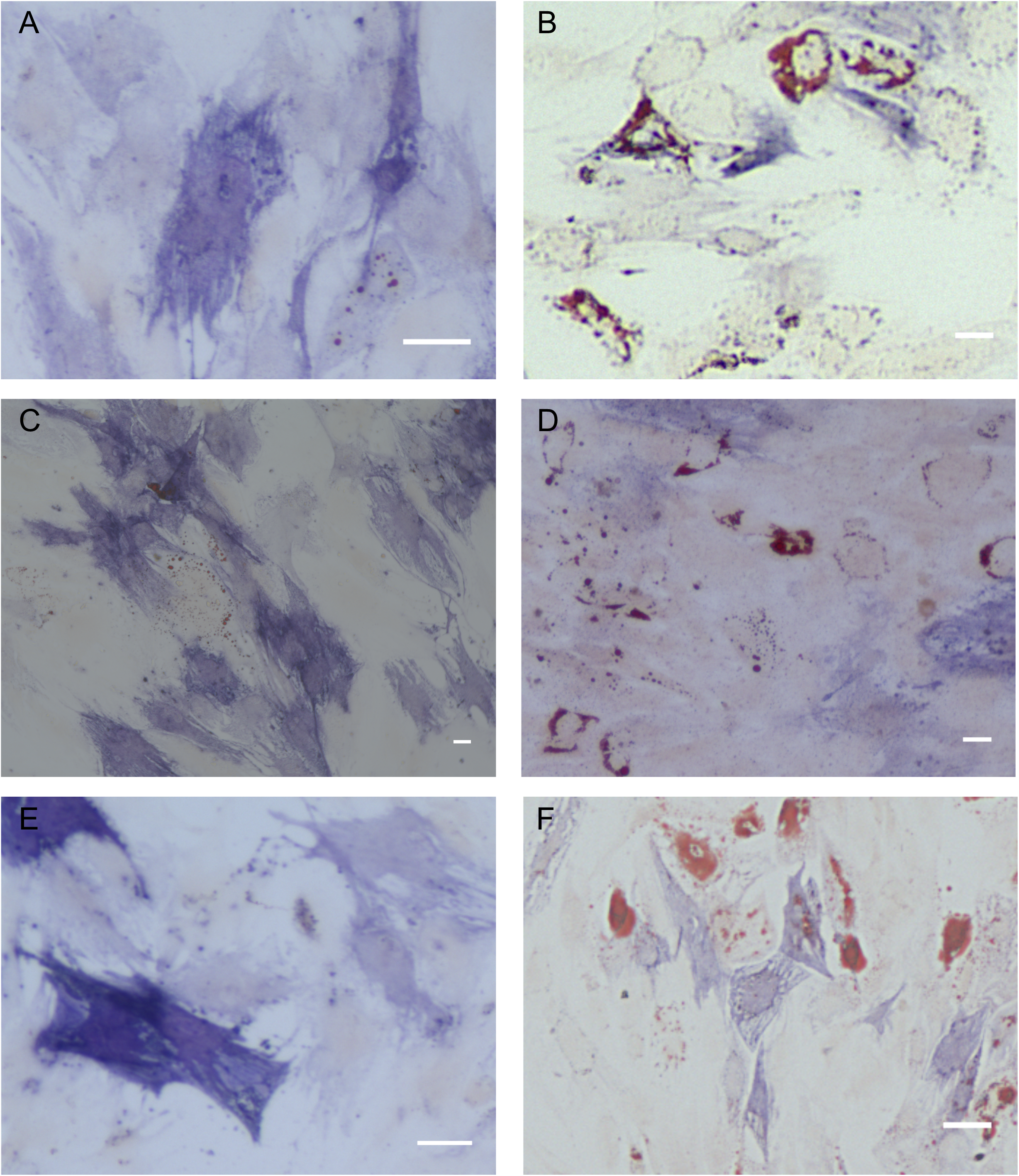
Differentiation experiments. ADSCs were cultured in mixed adipogenic and osteogenic differentiation media on each substrate for 14 days and then prepared for staining. ALP and lipid vacuoles were respectively stained by Fast blue RR and Oil red O on (A, C, E) non-porous and (B, D, F) 0.5 µm pore diameter SiO_2_ membranes (Scale bars = 50 µm).

